# ERO1α promotes hypoxic tumour progression and is associated with poor prognosis in pancreatic cancer

**DOI:** 10.1101/662411

**Authors:** Nikhil Gupta, Jung Eun Park, Wilford Tse, Jee Keem Low, Oi Lian Kon, Neil McCarthy, Siu Kwan Sze

## Abstract

Pancreatic cancer is a leading cause of mortality worldwide due to difficulty detecting early-stage disease and our poor understanding of the mediators that drive the progression of hypoxic solid tumours. We, therefore, used a heavy isotope ‘pulse/trace’ proteomic approach to determine how hypoxia alters pancreatic tumour expression of proteins that confer treatment resistance, promote metastasis, and suppress host immunity. Using this method, we identified that hypoxia stress stimulates pancreatic cancer cells to rapidly translate proteins that enhance metastasis (NOTCH2, NCS1, CD151, NUSAP1), treatment resistant (ABCB6), immune suppression (NFIL3,WDR4), angiogenesis (ANGPT4, ERO1α, FOS), alter cell metabolic activity (HK2, ENO2), and mediate growth-promoting cytokine responses (CLK3, ANGPTL4). Database mining confirmed that elevated gene expression of these hypoxia-induced mediators is significantly associated with poor patient survival in various stages of pancreatic cancer. Among these proteins, the oxidoreductase enzyme ERO1α was highly sensitive to induction by hypoxia stress across a range of different pancreatic cancer cell lines and was associated with particularly poor prognosis in human patients. Consistent with these data, genetic deletion of ERO1α substantially reduced growth rates and colony formation in pancreatic cancer cells when assessed in a series of functional assays *in vitro*. Accordingly, when transferred into a mouse xenograft model, ERO1α-deficient tumour cells exhibited severe growth restriction and negligible disease progression *in vivo*. Together, these data indicate that ERO1α is potential prognostic biomarker and novel drug target for pancreatic cancer therapy.

## Introduction

Pancreatic cancer is associated with <10% patient survival within just 5-years of diagnosis, reflecting a mortality rate approximately double that of other major cancer types (1-6). Pancreatic Ductal Adenocarcinoma (PDA) is the most common form of pancreatic malignancy but is typically diagnosed only in late-stage disease, hence both the incidence and deaths attributable to PDA continue to increase (7, 8). At present, there is no clinical procedure that can accurately detect early asymptomatic PDA, largely due to the lack of specific biomarkers of disease (9, 10), hence there is an urgent need for better predictors of tumour development and progression in this patient group with extremely poor prognosis.

Hypoxia frequently affects solid tumours that outgrow their local supplies of oxygen and nutrients. Previous studies have reported that the average pO_2_ of a developing tumour is just 0-5.3 mmHg, whereas the pO_2_ of adjacent healthy tissues ranges between 9.2 and 92.7 mmHg (11), indicating that hypoxia is a major influence on the biology of developing cancers. In particular, hypoxia-inducible factor (HIF-1α) is a key regulator of cellular responses to low-oxygen stress and appears to play a critical role in mediating tumour survival and rate of progression (12-15). In part, this is achieved via HIF-1α induction of pro-angiogenic mediators such as vascular endothelial growth factor (VEGF) (16-18), and transcription factors including Twist, Snail, and ZEB1 that promote epithelial-mesenchymal transition (EMT) (19, 20). Together, these effects significantly enhance neo-vascularisation of the tumour site, promote tissue invasion/metastasis, and increase chemotherapy resistance in many epithelial cancers. In pancreatic tumours, HIF-1α expression in a CD133+ stem cell-like population has been shown to promote EMT (21), and mutations in HIF-1α itself are key drivers of a variety of cancer types including PDA (22). These data are consistent with the emerging consensus that hypoxia-induced metabolic reprogramming is a hallmark feature of solid tumours including PDA (23), a key event driving the epigenetic changes that promote early invasion and metastasis (24-26), and a crucial determinant of immunosuppressive phenotypes that limit the effectiveness of many cancer therapies (27, 28). Despite substantial research progress, the molecular mechanisms that underpin hypoxia-induced effects on tumour development remain poorly understood, hence pancreatic cancer prognosis has failed to improve significantly for many years and treatment options for this disease remain extremely limited.

In the current study, we sought to determine the molecular basis of hypoxia effects on pancreatic cancer progression by using a pSILAC proteomic method (pulsed Stable Isotope Labelling of Amino acid in Cell culture) which facilitates analysis of how environmental factors impact on *de novo* protein synthesis via LC-MS/MS-based quantitation (29, 30). Using this approach, we investigated how the repertoire of proteins being actively translated by PDA cells is modified by nutrient starvation and hypoxia stresses, before comparing these data with patient survival statistics and pancreatic cancer gene expression profiles in publically available databases (PRECOG and GEO). These analyses revealed that the oxidoreductase enzyme ERO1α is actively translated by PDA tumours under both hypoxic and serum-free conditions, while also constituting a highly expressed gene associated with poor patient survival in both PRECOG and GEO datasets. Accordingly, genetic deletion of ERO1α inhibited pancreatic tumour proliferation, colony formation, and cellular ROS production *in vitro*, while cell transfer into a mouse xenograft model confirmed a profound reduction in tumour development *in vivo*. Together, these data indicate that hypoxia-induced enzyme ERO1α is a novel therapeutic target in pancreatic cancer and a potential biomarker for enabling earlier disease detection and better prediction of clinical course in human patients.

### Experimental methods

#### Cell Culture

Human pancreatic cells (MIAPaCa-2) were purchased from the American Type Culture Collection (Manassas, VA, USA) and cultured in Dulbecco’s Modified Eagle’s Medium (DMEM) (Biowest, France) supplemented with 10% foetal bovine serum (FBS) (Gibco, USA) and 1 % Penicillin/Streptomycin (Naclai-Teqsue, Japan) in a humidified incubator at 37°C with 5% CO_2_. For serum starvation experiments, the cells were washed three times in 1×PBS before incubating in FBS-free media for 24h. For hypoxic experiments, cells were subjected to low-oxygen culture (<0.1% O_2_, 5% CO_2_, 95% N_2_) in a hypoxia chamber for 24h. Cell lysates were subsequently obtained for protein and gene expression analyses. Cell proliferation was measured using MTT assays as previously described (31).

#### pSILAC labelling and proteomic sample preparation

MIAPaCa-2 cells were first grown for 24h in ‘light’ culture medium (containing unlabelled 146 mg/l ^12^C_6_, ^14^N_2_-L-Lysine and 84mg/l ^12^C_6_, ^14^N_2_-L-Arginine), then switched onto ‘heavy’ medium (containing labelled 146 mg/l ^13^C_6_, ^15^N_2_-D-Lysine and 84mg/l ^13^C_6_, ^15^N_2_-D-Arginine) and cultured for 24h under normoxic or hypoxic conditions in the presence or absence of FBS (n=2 independent biological replicates). Cell were then washed with cold PBS and lysed using 8M Urea buffer containing cOmplete™ EASYpack protease inhibitor cocktail (Sigma Aldrich, USA). In-solution digestion was performed as previously described (26). Extracted peptides were subjected to fractionation on an XBridge™ BEH C18 column (4.6 ×250 mm; Waters Corporation, Milford, MA, USA) and analysed by liquid chromatography-tandem mass spectrometry (LC-MS/MS).

Peptides were separated and analysed on a Dionex Ultimate 3000 RSLCnano system coupled to a Q-Exactive Hybrid Quadrupole-Orbitrap mass spectrometer (Thermo Fisher Scientific Inc, Germany). Separation was performed on a Dionex EASY-Spray 75µm×10 cm column packed with PepMap C18 3µm, 100 A° (ThermoFisher Scientific Inc. Germany) using solvent A (0.1% formic acid in 99.9% water) and solvent B (0.1% formic acid in 90% ACN) at flow rate of 300nl/min with a 60min gradient. Peptides were subsequently analysed using the Q-exactive MS with an EASY nanospray source (Thermo Fisher Scientific Inc, Germany) at an electrospray potential of 1.5kV. A full MS scan (350 – 1,600 m/z range) was acquired at a resolution of 70,000 at m/z 200 and a maximum ion accumulation time of 100ms. Dynamic exclusion was set as 15s. The resolution of high energy collisional dissociation (HCD) spectra was set to 17,500 at m/z 200. The automatic gain control settings of the full MS and MS2 scans were 3E6 and 2E5, respectively. The 10 most intense ions above a 2,000-count threshold were selected for HCD fragmentation, with a maximum ion accumulation time of 100ms. An isolation width of 2 was used for MS2. Single and unassigned charged ions were excluded from MS/MS. For HCD, the normalized collision energy was set to 28%. The underfill ratio was defined as 0.2%.

Protein identification and quantitation were performed by processing the raw data from three replicates using Protein Discoverer (PD) version 2.2 software (Thermo Scientific Inc. Germany). The MS/MS spectra were deisotoped and deconvoluted using the MS2 spectrum processor node in PD. Mascot and Sequest HT were used in parallel and the data compared against a protein sequence file from the UniProt human database (downloaded on 06 Feb 2017, 1,586,248 sequences, 61,972,042 residues). For searches using both engines, maximum missed cleavage sites per protein was set at 2, with precursor and fragment ions mass tolerance set at 10ppm and 0.02Da respectively. Carbamidomethylation (C) was set as a fixed/static modification. SILAC_R6 (R)/13C(6), SILAC_K8 (K)/13C(6)15N(2), acetylation (Protein N-term), deamidation (NQ) and Oxidation (M) were set as dynamic/variable modifications in both searches. A protein group list was generated from PD 2.2 software and the list was filtered to exclude contaminating proteins. Only proteins with q-value <0.01 (<1% FDR) as determined by percolator and detected in both replicates were used for further analysis. The mass spectrometry proteomics data have been deposited to the ProteomeXchange Consortium via the PRIDE partner repository with the dataset identifier PXD014087.

#### Western blot

Cells were washed twice in cold PBS and lysed in RIPA buffer (0.1% SDS, 1% NP-40, 1% sodium deoxycholate, 150mM NaCl, 1mM EDTA, and 50Mm Tris-HCl pH 7.4, with cOmplete™ EASYpack protease and phosphatase inhibitors) (Sigma-Aldrich, USA). Cell lysates were subjected to western blotting using the following primary antibodies: anti-ERO1α (CST), anti-HIF-1α (CST), anti-PDL1 (Abcam), anti-NFIL3 (CST), anti-PHD2 (CST), and anti-E-CAD (BD Bioscience). The loading control was beta-actin (1:5000) (EMD Millipore, USA). Proteins bound by each antibody were visualized using SuperSignal™ West Pico Chemiluminescent substrate (Thermo Fisher Scientific, USA).

#### RNA Isolation and qPCR

Total RNA was isolated using Nucleospin RNA II kits (Macherey-Nagel GmbH, Germany). Quantitative PCR (qPCR) reactions were performed using the CFX Connect™ Real-Time PCR Detection system (Bio-Rad Laboratories, Inc.) with KAPA SYBR® FAST qPCR Master mix. Actin was used as an internal control. The primer sequences used for qPCR are provided in Supplementary Table 1.

#### Bioinformatic Analysis

The pSILAC protocol allows newly synthesized proteins incorporating heavy (H) arginine and lysine to be distinguished from pre-existing proteins that include only light (L) arginine and lysine residues. Change in pSILAC H/L ratio under low-oxygen conditions can, therefore, be used to determine the impact of hypoxia on tumour cell protein expression activity. Proteins that were actively translated in response to hypoxia either in the presence of serum (HS/NS >1.5) or under serum-free conditions (HSF/NSF >1.5) (p<=0.001) were shortlisted for further analysis. Gene ontology and pathway analyses were performed using Functional Annotation (FunRich) (32). Prediction of Clinical Outcomes from Genomic profiles (PRECOG) and Gene Expression Omnibus (GEO) web tools were used to correlate the pSILAC proteomic data with clinical pancreatic tumour gene expression profiles in order to identify hypoxia-inducible proteins associated with poor prognosis (33, 34). Hypoxia-induced proteins that also displayed upregulated gene expression in cancer cells as reported in the GEO database were identified as potential targets. These targets were then further analysed using PRECOG to query associations between gene expression profiles and real-world patient outcomes. Potential targets associated with poor prognosis were identified by PRECOG meta z score. ERO1α displayed one of the highest meta z scores in the dataset and was therefore followed-up in subsequent functional studies. Datasets GSE15471, GSE78229, GSE67549 and GSE28735 reporting the results of pancreatic cancer microarrays in the NCBI GEO database (http://www.ncbi.nml.nih.gov/geo/) were analysed to confirm differential ERO1α expression (35-37).

#### CRISPR/Cas9 deletion of ERO1α

All CRISPR gRNA were computationally designed using CHOPCHOP V.2 software (Harvard, USA) to calculate ‘on target’ score and predicted Cas9 cleavage efficiency using a range of different algorithms (Benchling, San Francisco, CA, USA). Three gRNA were designed (Supplementary Table 2), and target sequences were cloned into lentiCRISPR v2 plasmids (Addgene plasmid # 52961) as described previously (38).

#### Colony formation and wound-healing assays

Colony formation assays were performed as previously described using a total of 200 cells/chamber seeded into six-well plates and cultured in DMEM (39). To quantify migratory potential, cells were spread and cultured until confluence, then scratched using a 200µl pipette tip before washing and incubating in fresh serum-free medium. After 24h culture, scratch dimensions were analysed using an optical microscope (40).

#### Intracellular ROS detection

Intracellular ROS analysis was performed as previously described (41). Briefly, cells were incubated with 10µM DCFDA in culture media for 1h at 30°C before harvesting, then washed twice in ice-cold PBS and examined under a fluorescent microscope (λex = 495 nm λem = 530nm) to identify green fluorescence generated by ROS production.

#### Xenografted tumour development assay

All animal studies were approved and conducted in compliance with the guidelines of the Institutional Animal Care and Use Committee (IACUC) of Nanyang Technological University. Ncr nude mice (males aged 7 weeks) were obtained from InVivos Pte, Ltd (Singapore). For tumour formation studies, mice were administered 3×10^6^ MIAPaCa-2-Con cells (n=6) or MIAPaCa-2-ERO1α^-/-^ (KO) cells injected subcutaneously into the neck area and tumour size was monitored by caliper measurement. Mice were euthanized when tumour size exceeded 2000mm^3^.

#### Statistical analysis

Statistical analysis performed using GraphPad Prism 6.0 software (GraphPad Softwares Inc., La Jolla, CA, USA). Statistical differences between variables were determined using Student’s paired t-test, and P<0.05 was considered to represent a statistically difference. FunRich tool provided p value of each ontology enrichment score. GEO datasets and PRECOG tool provide statistical analysis of overall survival and z scores for the high and low risk groups.

## Results

### Hypoxia induces dynamic changes in pancreatic cancer cell translational activity

The molecular mechanisms by which hypoxia favors pancreatic cancer progression and therapy resistance remain poorly understood. We hypothesized that hypoxic cancer cells co-opt normal metabolic pathways to force translation of proteins that support growth in unfavorable local conditions. To test this concept, we used a pSILAC proteomic approach to investigate how the pancreatic tumour cell proteome is modified in response to a hypoxic microenvironment. To mimic tumour hypoxia *in vivo*, we first cultured pancreatic ductal adenocarcinoma (PDA) cells under low-oxygen conditions *in vitro*, either in the presence or absence of serum, and then compared viability with cells maintained on normal oxygenation (normoxia) throughout. When assessed in MTT assays, PDA cells displayed stable growth rates over a 24h culture period irrespective of serum supplementation, and cell viability remained essentially unchanged thereafter. In contrast, PDA cells subjected to low-oxygen conditions displayed reduced proliferation after just 24h culture (Figure 1A), while substantially upregulating the characteristic hypoxia markers HIF-1α, hexokinase 2 (HK2) and N-Myc Downregulated protein1 (NDRG1) (Figure 1B). These data confirmed that our model system reproduces the oxygen-deprived state encountered by solid pancreatic tumours *in vivo* and replicates their associated cellular responses *in vitro* (42, 43). We, therefore, proceeded to culture PDA cells with light-medium (^12^C_6_, ^14^N_2_-L-Lysine, ^12^C_6_, ^14^N_2_-L-Arginine) before switching to heavy medium (^13^C_6_, ^15^N_2_-D-Lysine, ^13^C_6_, ^15^N_2_-D-Arginine) and subjecting the cells to 24h culture under normoxia or hypoxia in the presence or absence of serum. Upon completion of culture, cellular proteins were extracted and analysed by LC-MS/MS in order to identify molecules that incorporated heavy isotope-labelled Lys and Arg (synthesized *de novo* in response to hypoxia stress) and differentiate these from ‘light’ proteins from the pre-existing pool within the tumour cells (n=2 biological replicates; Supplementary Data 1). Adjusted p values were generated from PD 2.2 software for pSILAC analysis. We observed that PDA cells expressed a wide range of different proteins after culture, whether conducted in normoxia under serum replete (5684) or serum-free conditions (5944), or following hypoxia with serum provided (5932), or withheld for the duration (5677) (Figure 1C). Unsurprisingly, the majority of these proteins were derived from a common pool of shared molecules (n=5010), but a substantial fraction were specifically modulated in response to changing microenvironmental conditions (threshold >0.6 log_2_ Fold Change in H/L ratio) (Figure 1D). These candidate mediators of the tumour response to hypoxia stress were subsequently screened and cross-validated by RT-qPCR and western blot in order to verify the pSILAC data. Distribution analyses revealed that most of these proteins were substantially downregulated during hypoxia stress, regardless of serum provision, including several key mediators of cell translation, metabolism, and mRNA maturation processes (Supplementary Figure 1).

**Figure 1.**
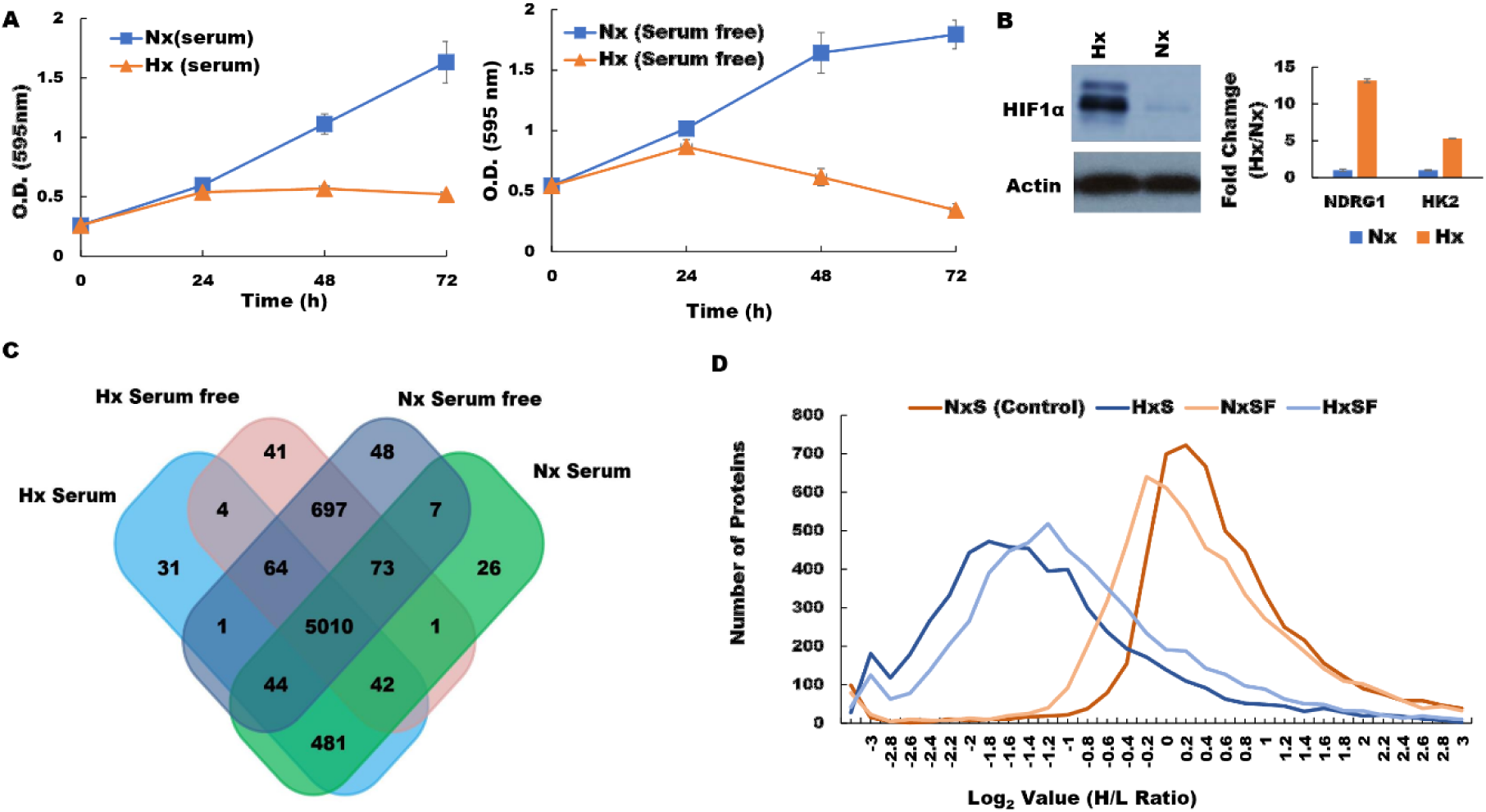
Hypoxia modifies the pancreatic cancer cell proteome under both serum-replete (S) and serum-free conditions (SF). **(A)** PDA cell proliferation curve during culture in S and SF conditions confirming growth restriction during hypoxia. **(B)** Western blot of HIF-1α protein and RT-PCR assessment of fold change (Hx/Nx) in hypoxia markers NDRG1 and HK2. **(C)** number of proteins identified in PDA cells after culture under the different conditions indicated. **(D)** Analysis of protein distribution against log_2_ H/L ratio for the S and SF culture conditions during normoxia/hypoxia revealed that the majority of these proteins are downregulated during hypoxia.

Among the proteins induced by pancreatic tumour hypoxia (high Hx/Nx ratios), we observed that 206 were actively synthesized under serum-free conditions, 138 were translated in the presence of serum, and 20 were actively expressed in both conditions (Figure 2A). Comparisons between these groups based on functional classification revealed that the majority of hypoxia-sensitive proteins were mediators of signalling pathways/communication and cell growth, but we also detected marked hypoxia effects on key regulators of protein and nucleic acid metabolism. Furthermore, proteins (NOTCH2, NCS1, NUSAP1,CD151) that are known to promote metastasis were increased more than 10 times under hypoxia indicates how hypoxia can change the tumor dynamics. Additionally, some of the proteins confers to immune suppression (NFIL3, WDR4) and drug resistance (ABCB6) were also increased under hypoxia may contribute to the poor survival of PDA patients. Further analysis indicated that several critical mediators of cellular transport and metabolic activity were more potently induced by hypoxia under serum-free conditions (Figure 2B & Supplementary data 2), indicating that nutrient availability plays a major role in determining how oxygen restriction modifies pancreatic tumour biology. Among the 20 proteins induced by hypoxia irrespective of serum presence/absence were several well-established mediators of cancer progression, including key oncogenic transcription factors FOS and JUN, as well as the pro-angiogenic factor ANGPTL4 which is directly involved in neo-vascularization events (Figure 2C). Several other proteins in this group also displayed greater induction under serum-free conditions (NOL3, NCS1, CD151), suggesting synergistic effects of hypoxia and nutrient deprivation on pancreatic cancer cell development and angiogenic activity (Figure 2D). To confirm these findings, gene expression levels were validated by quantitative PCR analyses, which were highly consistent with the pSILAC proteomic data (see Supplementary Figure 2). Having identified a range of hypoxia-induced proteins that are expressed by pancreatic cancer cells *in vitro*, it remained unclear which of these mediators exerted the greatest impact on malignant disease progression *in vivo*. We, therefore, proceeded to investigate whether hypoxia-induced changes in pancreatic cancer cell proteome impact on disease progression and modify patient outcome using real-world clinical data.

**Figure 2.**
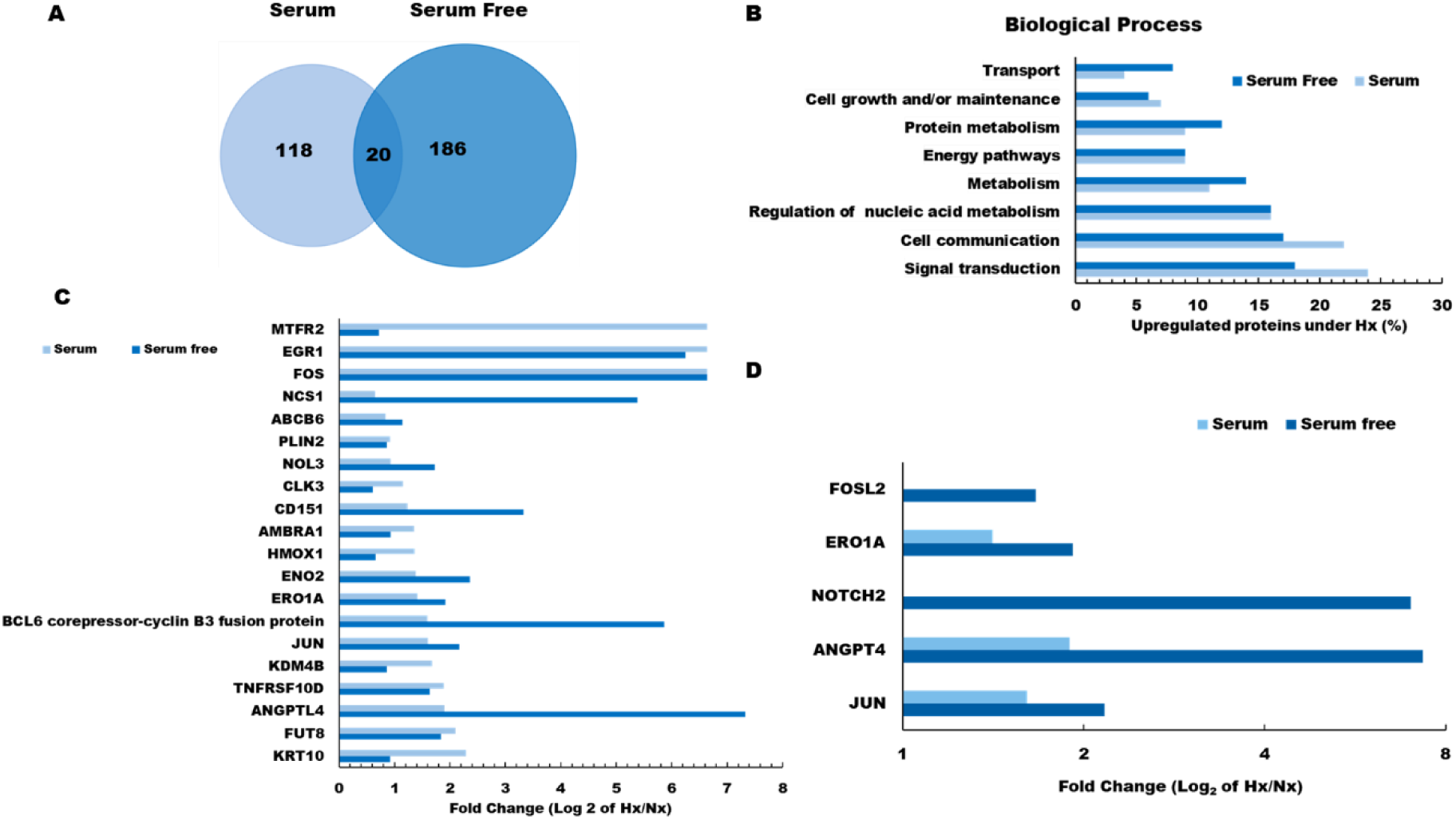
Gene ontology analysis of hypoxia effects on the pancreatic tumour cell proteome: **(A)** Venn diagram showing the number of hypoxia-induced proteins identified as being newly synthesized under S or SF conditions. **(B)** Functional classification of tumour proteins expressed *de novo* in response to hypoxia stress. **(C)** Log_2_ fold change (Hx/Nx) in actively translating proteins detected in either S or SF culture conditions, and **(D)** Log_2_ fold change (Hx/Nx) in angiogenic factor expression comparing S and SF culture conditions.

### Oxygen restriction induces pancreatic cancer cell expression of metabolic mediators that predict poor clinical outcome

To determine the clinical relevance of hypoxia-induced changes in protein expression by PDA cells, we conducted ‘PREdiction of clinical outcome from genomic profiles’ (PRECOG) analysis to identify possible correlations of our proteomic results with patient survival data.(28) Hypoxia-induced proteins were first integrated into PRECOG and meta z scores calculated to determine the confidence level of associations with adverse or favorable clinical outcomes (Figure 3A). Using this approach, we observed that several proteins associated with poor prognosis were more strongly induced by hypoxia during serum starvation (Figure 3B), and that meta z scores for the pancreatic cancer dataset were generally higher in serum-free relative to serum-replete conditions. Among these were several proteins for which high gene expression levels were strongly associated with poor patient prognosis, suggesting a likely role in hypoxia-driven PDA progression. In particular, the oxidoreductase protein ERO1α displayed an unweighted meta z score of 7.67 for all cancer types, thus strongly implicating this enzyme as a key determinant of poor clinical outcome in human pancreatic cancer. These data were then verified by qRT-PCR and western blot analyses, which confirmed that PDA cells express ERO1α at levels consistent our proteomic results (Figure 4A & 4B). Further analysis of ERO1α expression patterns in relevant GEO databases,(44) enabled us to confirm that a range of different pancreatic cancer cell lines strongly upregulates ERO1α under hypoxic conditions (Figure 3F). Mining of additional publicly available datasets (Supplementary Table 2) revealed that this enzyme is also expressed at significantly higher levels in pancreatic tumour samples compared with adjacent healthy tissue (Figure 3C & 3D), in clear association with adverse patient outcomes (Figure 3E). Together, these data indicate that under hypoxic conditions, pancreatic tumour cells significantly upregulate the enzyme ERO1α, which is a critical determinant of pancreatic cancer patient survival. While ERO1α plays a well-established role mediating disulfide bond formation in a range of different secreted and membrane proteins, the survival advantages this confers on PDA cells and the mechanisms involved remained unknown.

**Figure 3.**
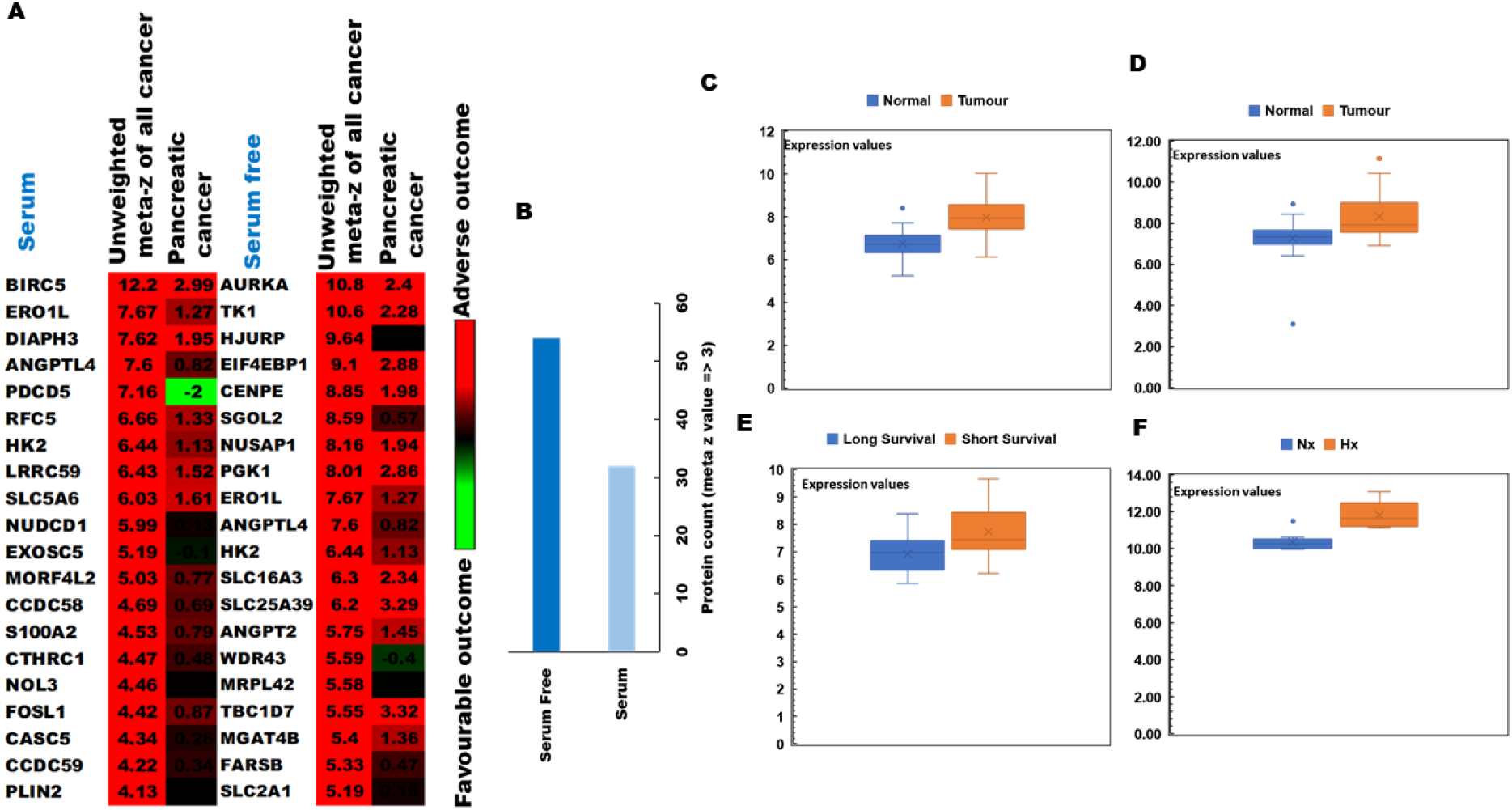
PRECOG analysis of the pancreatic tumour proteome and correlations with patient outcomes: **(A)** Top 20 genes associated with poor clinical prognosis in cancer patients and identified in our pSILAC dataset of hypoxia-induced proteins (high meta z values indicating proteins most strongly associated with adverse [red] or favourable [green] survival data) **(B)** Protein count in serum-replete (S) and serum-free (SF) conditions, indicating that markers associated with poor prognosis are more strongly induced by hypoxia (meta z value >3) during SF culture. **(C)** Comparison of ERO1α expression levels between healthy tissues and disease samples in relevant GEO datasets: GSE28735; 45 pancreatic tumour samples and 45 matching normal samples **(D)** GSE15471; 78 pancreatic cancer and normal tissue samples **(E)** GSE78229; 50 pancreatic tissue samples grouped by survival duration, and **(F)** GSE67549; n=9 pancreatic cancer cell lines either cultured in Nx or subjected to Hx. Database selection criteria are displayed together with an overview of the 4 studies selected in Supplementary Table 2.

**Figure 4.**
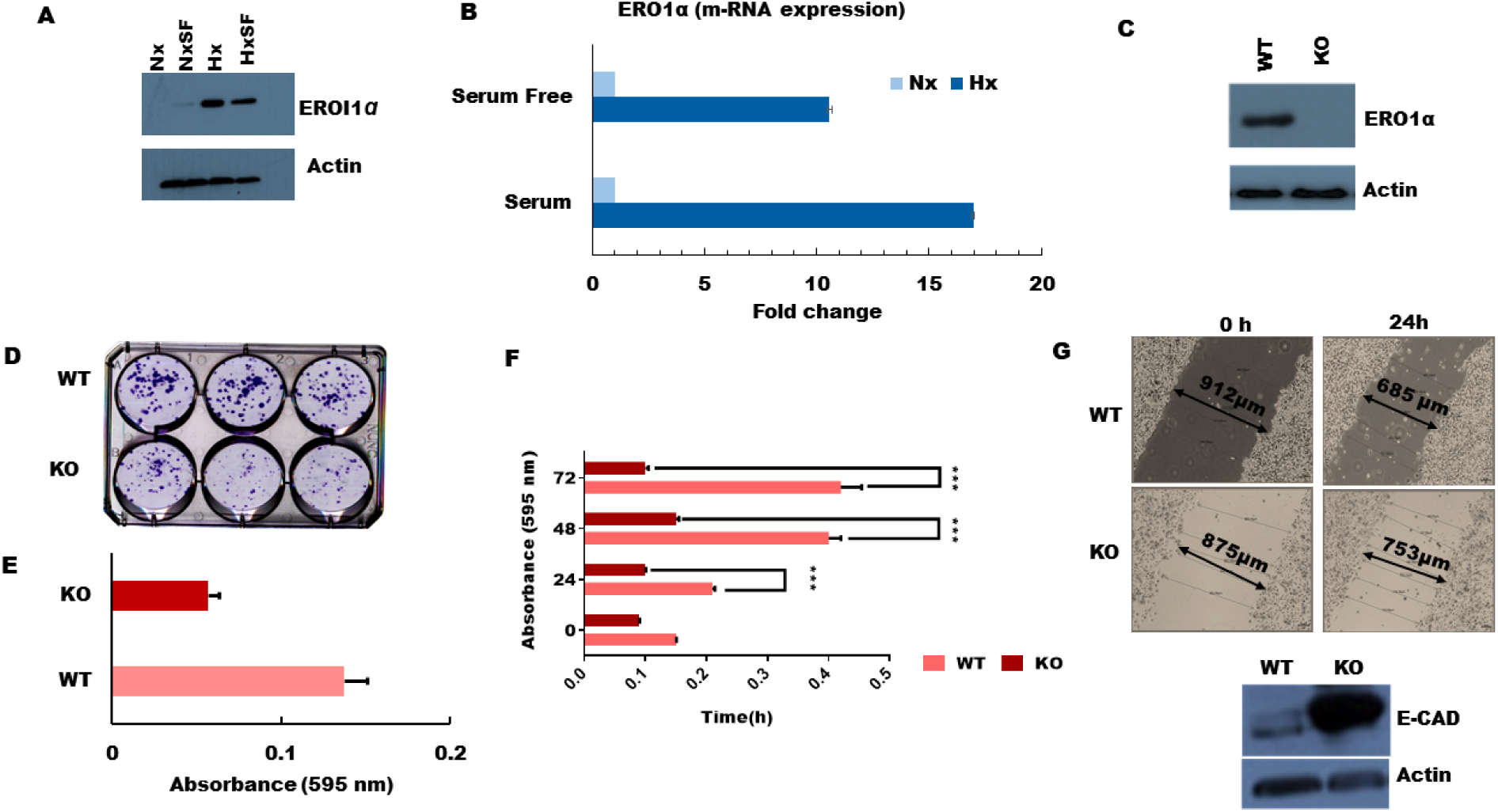
ERO1α deletion reduces PDA tumour cell growth, colony formation, and migratory potential *in vitro*: (**A)** Western blot confirming that ERO1α protein is highly expressed during tumour hypoxia in both S and SF conditions, consistent with the pSILAC data and patient gene expression profiles in the PRECOG and GEO datasets. (**B)** ERO1α mRNA expression levels as assessed using qPCR. **(C)** Western blot analysis of ERO1α expression in WT and ERO1α-KO PDA cells. **(D, E)** Colony formation potential of PDA cells comparing WT and ERO1α-KO clones. (**F)** Cell proliferation curves of WT and ERO1α-KO clones in hypoxic MTT assays. **(G)** Wound-healing assay performed using the indicated cell clones and culture conditions, demonstrating the reduced migratory potential of ERO1α-KO tumour cells. **(H)** Western blot analysis of the archetypal EMT suppressor protein E-cadherin, which was overexpressed in ERO1α-KO PDA cells. ***P<0.001.

### ERO1α mediates pancreatic cancer cell growth, ROS production, and tumorigenicity in vivo

To better understand the role of ERO1α in pancreatic cancer progression, we next deleted the corresponding gene using CRISPR/Cas genome editing technology and assessed the impact on tumour cell function both *in vitro* and *in vivo*. For this, we selected a sgRNA with the highest ‘on target’ score, which was directed against exon 7 of human ERO1α and predicted to induce a frameshift mutation that generates non-functional/truncated gene products. We then infected human pancreatic cancer cells with Cas9-expressing lentiviral vectors in order to disrupt the ERO1α gene, and used Sanger sequencing to verify mutation at the predicted site. Accordingly, western blot analysis of ERO1α expression in the treated cells confirmed that our gene disruption strategy achieved complete knockout of ERO1α protein (Figure 4C), so we proceeded to test whether pancreatic tumour cell proliferation rates differed between WT and ERO1α-KO clones. Unlike their WT counterparts, ERO1α-KO tumours displayed little to no growth over a 24-72h culture period (Figure 4F), and colony formation was reduced by as much as 50% (Figure 4D & 4E), suggesting that ERO1α deletion significantly reduces the replicative capacity of pancreatic cancer cells. Furthermore, when assessed in a series of wound healing assays, we observed that ERO1α-KO tumour cells displayed impaired ability to close the injury due to migration rates being reduced by >50% relative to ERO1α-competent tumour cells (Figure 4G). Given these marked defects in cell motility, we next assessed tumour cell expression of cadherins, which play a central role in restricting cancer cell migration and tissue invasion. When analysed by western blot, key EMT suppressor E-cadherin was found to be highly expressed in ERO1α-KO tumour cells, perhaps explaining their reduced migratory potential in our functional assays (Figure 4H).

Having detected a clear impact of ERO1α deletion on PDA cell expression of a key protein regulator of tumour development, we next investigated whether ERO1α was also required to induce other critical mediators of disease progression. In particular, HIF-1α is a key regulator of many oncogenic pathways known to be triggered by hypoxia, and our western blot analyses revealed a significant deficit in expression of this transcription factor in ERO1α-KO cells (Figure 5A) (45). Furthermore, in the absence of ERO1α, we observed that PDA cells also displayed substantially reduced expression of the immune inhibitory molecule PD-L1, which is typically upregulated in many different cancer types. (Figure 5A). Given that HIF-1α expression is known to be controlled by reactive oxygen species (ROS), we next assessed whether the generation of these mediators might represent a critical mechanism by which ERO1α influences pancreatic cancer progression. Indeed, previous work has identified that ERO1α oxidizes PDI which in-turn catalyzes the formation of disulfide bonds in folding proteins, which simultaneously generates intracellular hydrogen peroxide (H_2_O_2_) that can mutate DNA and trigger oncogenic events. When assessed in fluorescence based DCDFA assays, we observed that PDA cells lacking ERO1α displayed substantially reduced ability to generate ROS, exhibiting only 66% of the ROS capacity detected in WT tumour cells (Figure 5B & 5C). Together, these data confirmed that ERO1α-deficient PDA cells exhibit impaired tumour development *in vitro*, so we next performed xenograft experiments in Ncr-nude mice to determine whether these cells also displayed reduced oncogenic potential *in vivo*. When administered by subcutaneous (s.c) injection, ERO1α-competent tumours developed readily within just 2 weeks of cell transfer, whereas ERO1α-deficient cancer cells failed to drive tumour formation in recipient animals (n= 6 per group). Indeed, when the skin was excised from the injection sites we were unable to detect significant tumour development in any mice that had received ERO1α-KO PDA cells (Figure 5D and E). These data confirm that in the absence of ERO1α, pancreatic tumour cells undergo regression rather than progression *in vivo*.

**Figure 5.**
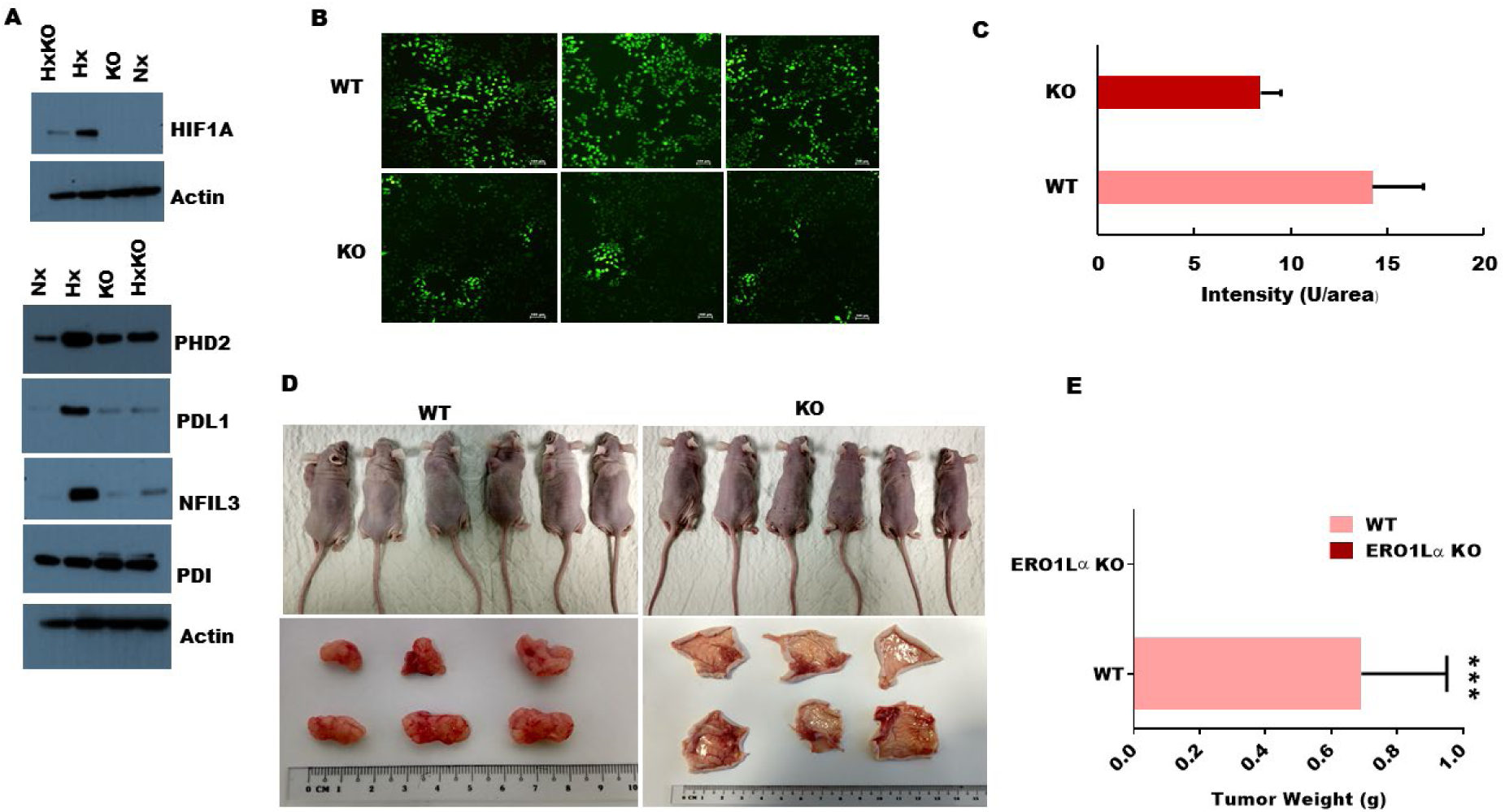
Impact of ERO1α deletion on oncogenic protein expression, ROS production, and xenograft tumour progression *in vivo*: **(A)** Western blot analysis of ERO1α-regulated proteins comparing WT with ERO1α-KO tumour cells. **(B)** DCFDA assay with representative fluorescence images of WT and ERO1α-KO PDA cells showing a significant reduction in ROS generation in ERO1α-deficient tumours. **(C)** Fluorescence intensity as measured by microplate reader confirming significant ROS impairment in ERO1α-KO clones (**D)** Representative images of xenografted mouse tumour injection sites before and 28 days after inoculation with the indicated cancer cells. **(E)** Tumours were excised and weighed; mean values are shown (n=6). ***P<.001 (student t-test).

## Discussion

Low oxygen availability in solid tumours is now well established as a crucial factor driving malignant progression, promoting treatment resistance, and conferring poor clinical outcome in pancreatic cancer.(46) Since the molecular basis of these effects remains poorly defined, we used a pSILAC proteomic approach to identify hypoxia-sensitive proteins expressed by pancreatic cancer cells when subjected to oxygen restriction *in vitro* either in the presence or absence of serum supplementation (mimicking restricted blood/nutrient supply *in vivo*). Using this model, we detected rapid induction of several established biomarkers of tumour hypoxia, and observed that active protein translation was broadly downregulated in the oxygen-deprived microenvironment. However, we also identified that hypoxia stress induces pancreatic tumour cell expression of the oxidoreductase protein ERO1α, which displayed a clear association with poor survival statistics mined from patient gene expression databases.

Hypoxia is known to impede the growth of both tumours and healthy cell types, but variable severity and duration of hypoxia can induce a range of different effects that have been shown to increase cancer cell viability, therapy resistance, and metastatic potential (45, 47, 48). While tumour translational activity was generally downregulated during low-oxygen stress in our assays, we also detected a group of proteins that were instead selectively induced by hypoxia (206 in serum-free cultures, 138 under serum-replete conditions, and 20 that were actively synthesized in both settings). The relatively high number of tumour proteins upregulated in serum-free culture might indicate translation of molecules that support cancer cell survival under adverse conditions. In support of this concept, the profile of proteins induced in the presence/absence of serum displayed divergent functional profiles, with serum-replete culture favouring mediators of cell communication and signal transduction, whereas serum-free conditions induced regulators of cellular transport and metabolic processes. Hypoxia induced several proteins have shown to increase metastasis, immune suppression and treatment resistant in different cancers. One such highly upregulated (Hx/Nx >10) protein in our dataset NCS1 (neuronal calcium sensor 1) a calcium binding protein involved in many cellular processes have shown to promote metastasis in breast cancers.(49) Furthermore, our data indicated that hypoxic tumours subjected to serum starvation displayed greater induction of the anti-apoptotic protein NOL3, a key mediator of tumour migration/invasion CD151, and angiogenic factor ANGPTL4 which supports tumour neovascularization. Together, these data indicate that hypoxia effects on pancreatic tumour development are strongly influenced by the nutrient status of the constituent cancer cells, which subsequently upregulate key mediators of cell survival, dissemination, and angiogenic processes.

In order to determine the clinical relevance of hypoxia-induced protein translational activity in pancreatic cancer cells, we integrated our data with cancer patient survival statistics available via public gene expression databases. Using the ‘PRECOG’ tool to identify associations between gene expression levels and clinical outcomes, we were able to identify which hypoxia-triggered proteins exert the greatest influence on cancer patient prognosis. Intriguingly, proteins associated with poor prognosis were more actively translated by hypoxic tumours in serum-free cultures than in serum-replete conditions, further indicating that nutrient availability is a major determinant of how hypoxia impacts cancer progression. Further screening of these proteins revealed that ERO1α was associated with a particularly poor prognosis and displayed a high meta z score irrespective of serum availability. We, therefore, analyzed the expression of ERO1α in relevant GEO datasets which confirmed that ERO1α is induced by hypoxia stress across a range of pancreatic cell lines. These analyses also indicated that ERO1α is expressed at elevated levels in pancreatic cancers relative to adjacent healthy tissues from the same patient, and that high tumour expression of this enzyme is significantly correlated with short survival duration.

Previous studies have implicated ERO1α in a range of different cancer types, with current evidence suggesting potential roles in cancer cell invasion of healthy tissues, acquisition of chemoresistant properties, and promotion of angiogenesis (50). Recent data have also suggested that ERO1α-mediated protein folding is required for the generation of myeloid-derived suppressor cells that protect tumours against immune destruction (51). In order to test the function of ERO1α in pancreatic cancer cells, we used a CRISPR/Cas9 approach to delete the corresponding gene and assessed the impact on the oncogenic potential in a range of different assays. While ERO1α-KO pancreatic cancer cells were morphologically indistinguishable from their wild-type counterparts, tumour proliferation and colony forming potential were significantly reduced in the absence of this enzyme. Similarly, when we assessed the motility of ERO1α-KO tumour cells using *in vitro* wound healing assays, we observed a marked reduction in migratory potential compared with WT clones. Since ERO1α enzyme activity is known to generate reactive oxygen species (ROS) which directly regulate the hypoxia response through transcription factor HIF-1α, we next assessed whether ROS generation might underpin the influence of ERO1α on pancreatic cancer progression. Using DCFDA assays to quantify ROS levels in hypoxic tumour cells, we observed that ERO1α-KO clones displayed a significant reduction in ROS generation relative to WT clones. Accordingly, when WT or ERO1α-KO cancer cells were injected into BALB/c nu/nu mice, the enzyme-deficient tumours failed to develop whereas wild-type cells were highly tumorigenic *in vivo*.

Taken together, these data indicate that a combination of microenvironmental hypoxia and restricted blood/nutrient supply trigger pancreatic cancer cell translation of a range of different proteins that act to increase malignancy and are associated with poor clinical outcome. In particular, the oxidizing enzyme ERO1α is predominantly activated only under abnormal conditions such as hypoxia, which perhaps explains the marked induction of this protein in oxygen-starved tumours. Consistent with this concept, our data confirmed that ERO1α-deficient pancreatic cancer cells displayed negligible ability to form tumours *in vivo*, thus implicating ERO1α as a potential novel target for cancer therapy in human patients. Given that ERO1α function appears less critical under steady-state conditions, it is also possible that therapeutic disruption of this enzyme may succeed in limiting tumour growth and metastasis without significantly damaging nearby healthy cells and tissues. Further investigations will now be required to assess this possibility. Collectively, the findings of this systematic study using pSILAC-based quantitative proteomics reveal that tumour hypoxia induces active translation of a range of critical proteins that are associated with poor prognosis in human cancer patients. Specifically, our data reveal that the oxidoreductase enzyme ERO1α is a major proliferation regulator in pancreatic cancer cells both *in vitro* and *in vivo*, hence this protein may constitute a useful diagnostic marker of tumour progression and possible new therapeutic target in pancreatic malignancy.

## Data availability

LC-MS/MS raw data from the pulsed-SILAC experiments and results for protein and peptide identification and quantification from PD2.2 have been deposited to the ProteomeXchange Consortium via the PRIDE(52) partner repository with the dataset identifier PXD014087. The raw data and search results can be accessed from the PRIDE data depository. **Project Name:** Proteomics analysis of pancreatic cancer cell line (MiaPaCa-2)

## ACKNOWLEDGEMENTS

This work is in part supported by the Singapore Ministry of Education (MOE2016-T2-2-018 and MOE2018-T1-001-078) and Singapore National Medical Research Council (NMRC/OFIRG/0003/2016)

