## Supplementary Table and Figures for "ERO1α promotes hypoxic tumour progression and is associated with poor prognosis in pancreatic cancer"

**Hypoxia sensitive ERO1**α **promotes pancreatic cancer progression and associated with poor clinical prognosis**

**Supplementary Materials**

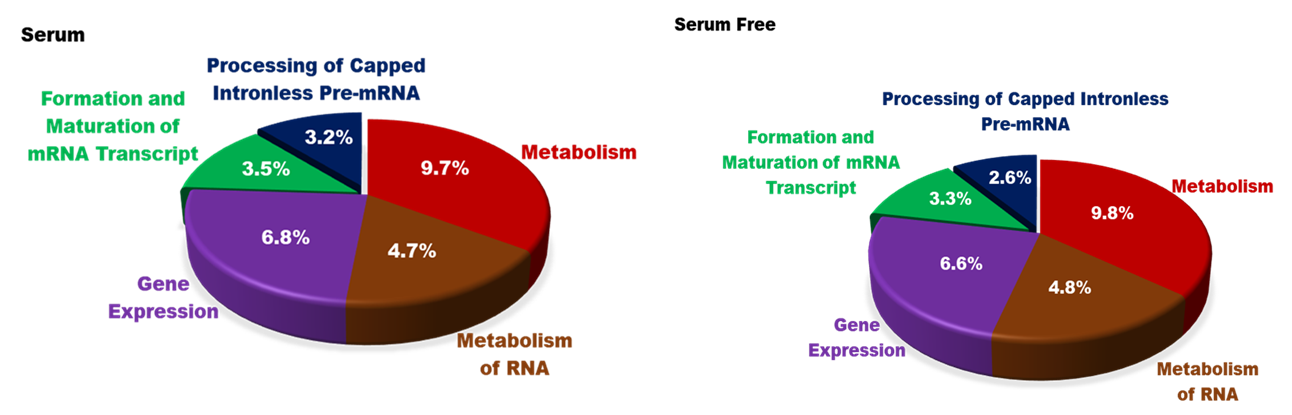

**Supplementary figure 1:** **Gene ontology analysis of the proteins downregulated by hypoxic tumour cells in either serum-replete or serum-free conditions.**

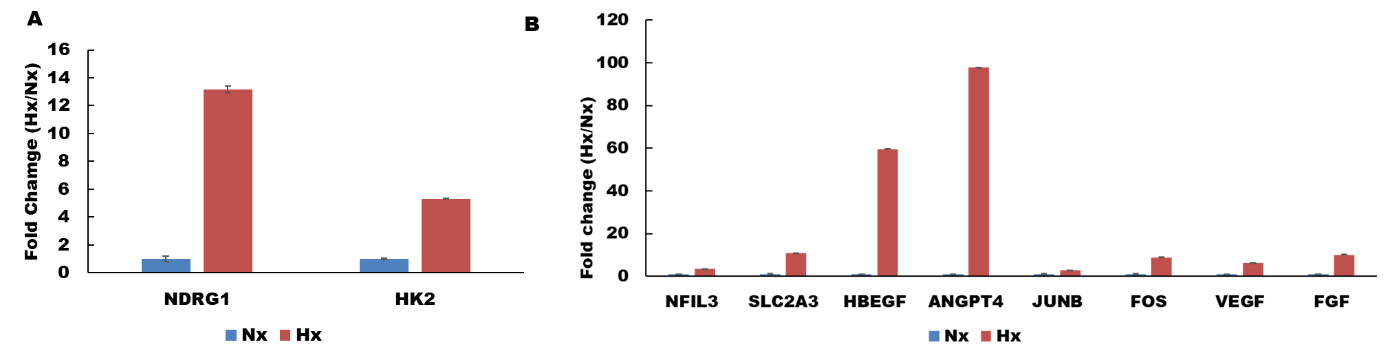

**Supplementary figure 2: qPCR validation of key hypoxia-induced proteins as identified by pSILAC analysis (expressed as Hx/Nx fold-change in gene expression level).**

**Supplementary Table 1 Primer sequences for quantitative PCR**

| **Primer** | | **5' - 3'** |
| --- | --- | --- |
| ERO1α_FW | ATGACATCAGCCAGTGTGGA | |
| ERO1α_RV | CATGCTTGGTCCACTGAAGA | |
| NFIL3_FW | AATGCAGACCGTCAAAAAGG | |
| NFIL3_RV | TTGTTCTTCCCCACACTTCC | |
| HBEGF_FW | GGTGGTGCTGAAGCTCTTTC | |
| HBEGF_RV | GCTTGTGGCTTGGAGGATAA | |
| VEGFA_FW | AAGGAGGAGGGCAGAATCAT | |
| VEGFA_RV | ATCTGCATGGTGATGTTGGA | |
| FGF1_FW | TGCCTCCAGGGAATTACAAG | |
| FGF1_RV | TATAAAAGCCCGTCGGTGTC | |
| ACTIN_FW | GGACTTCGAGCAAGAGATGG | |
| ACTIN_RV | AGCACTGTGTTGGCGTACAG | |
| SLC2A3_FW | ACCGGCTTCCTCATTACCTT | |
| SLC2A3_RV | AGGCTCGATGCTGTTCATCT | |
| ANGPTL4_FW | GCCTATAGCCTGCAGCTCAC | |
| ANGPTL4_RV | AGTACTGGCCGTTGAGGTTG | |
| JUNB_FW | AATGGAGGACACGGACTTTG | |
| JUNB_RV | ATGTCTCTGGCATCCCTACG | |
| FOS_FW | AGAATCCGAAGGGAAAGGAA | |
| FOS_RV | CTTCTCCTTCAGCAGGTTGG | |
| NDRG1_FW | ACAACCCTGAGATGGTGGAG | |
| NDRG1_RV | TGTGGACCACTTCCACGTTA | |
| HK2_FW | TCTATGCCATCCCTGAGGAC | |
| HK2_RV | TCTCTGCCTTCCACTCCACT | |

**Supplementary Table 2: List of gene expression microarray dataset used for ERO1Lα expression analysis**

| **S.NO.** | **GEO accession** | **Platform** | **Sample** | **Cancer Type** |
| --- | --- | --- | --- | --- |
| 1 | GSE15471 | GPL570 | Whole-tissue pancreatic ductal adenocarcinoma | Pancreatic Cancer |
| 2 | GSE78229 | GPL6244 | Pancreatic tumours tissue | Pancreatic Cancer |
| 3 | GSE67549 | GPL15207 | Cancer cell line | Pancreatic Cancer |
| 4 | GSE28735 | GPL6244 | Pancreatic tissue and adjacent normal tissue | Pancreatic Cancer |

**Supplementary Table 3**

| **ERO1Lα CRIPSR oligo design** | |  |  |
| --- | --- | --- | --- |
| **Name** | **Sequence** | **Scale** | **Purification** |
| **Exon 7 CHOP CHOP (No mismatch)** |  |  |  |
| ERO1Lα_Oligo1_LENTI_guide | CACCGGAGCGCTACACTGGTTACAA | 25nm | STD |
| ERO1Lα_Oligo2_LENTI_guide | AAACTTGTAACCAGTGTAGCGCTAC | 25nm | STD |
| **Exon 1 CHOP CHOP (3 mismatch)** | **Low Efficiency** |  |  |
| ERO1Lα2_Oligo1_LENTI_guide | CACCGCGCGGCTGGGGATTCTTGTT | 25nm | STD |
| ERO1Lα2_Oligo2_LENTI_guide | AAACAACAAGAATCCCCAGCCGCGC | 25nm | STD |
| **Exon 1 CHOP CHOP (3 mismatch)** | **High Efficiency** |  |  |
| ERO1Lα3_Oligo1_LENTI_guide | CACCGGCCGCGGCCCATTGCAGCTC | 25nm | STD |
| ERO1Lα3_Oligo2_LENTI_guide | AAACGAGCTGCAATGGGCCGCGGCC | 25nm | STD |

**Supplementary Data 1**

See Supplementary File 1.

**Supplementary Data 2**

See Supplementary File 2.
